# Controlling the switch from neurogenesis to pluripotency during marmoset monkey somatic cell reprogramming with self-replicating mRNAs and small molecules

**DOI:** 10.1101/2020.05.21.107862

**Authors:** Stoyan Petkov, Ralf Dressel, Ignacio Rodriguez-Polo, Rüdiger Behr

**Affiliations:** Platform Degenerative Diseases, German Primate Center GmbH, Leibniz Institute for Primate Research, Göttingen, Germany; Institute for Cellular and Molecular Immunology, University Medical Center Göttingen, Göttingen, Germany; German Center for Cardiovascular Research (DZHK), partner site Göttingen, Germany

## Abstract

Induced pluripotent stem cells (iPSCs) hold enormous potential for the development of cell-based therapies for many currently incurable diseases. However, the safety and efficacy of potential iPSC-based treatments need to be verified in relevant animal disease models before their application in the clinic. Moreover, in order to reduce possible risks for the patients, it is necessary to use reprogramming approaches that ensure to the greatest extent possible the genomic integrity of the cells. Here, we report the derivation of iPSCs from common marmoset monkeys (*Callithrix jacchus*) using self-replicating mRNA vectors based on the Venezuelan equine encephalitis virus (VEE-mRNAs). By transfection of marmoset fetal fibroblasts with Tomato-modified VEE-mRNAs carrying the human *OCT4, KLF4, SOX2*, and *c-MYC* (VEE-OKS-iM-iTomato) and culture in medium supplemented with two small molecule inhibitors, we first established intermediate primary colonies with neural progenitor-like properties. In the second reprogramming step, we converted these colonies into transgene-free pluripotent stem cells by further culturing them with customized marmoset iPSC medium in feeder-free conditions. The resulting cell lines possess pluripotency characteristics, such as expression of various pluripotency markers, long-term self-renewal, stable karyotype, and ability to differentiate into derivatives of the three primary germ layers *in vitro* and *in vivo*. Our experiments reveal a novel paradigm for flexible reprogramming of somatic cells, where primary colonies obtained by a single VEE-mRNA transfection can be directed either towards the neural lineage or further reprogrammed to pluripotency. These results (i) will further enhance the role of the common marmoset as animal disease model for preclinical testing of iPSC-based therapies and (ii) establish an in vitro system to experimentally address developmental signal transduction pathways in primates.

## INTRODUCTION

Despite significant advances in modern medicine, many diseases that increase the mortality or cause disabilities in the world’s human population currently remain incurable. Due to their properties that allow them to be expanded indefinitely *in vitro* and to be differentiated into various somatic cell types, pluripotent stem cells (PSCs) hold significant potential for the development of cell-based therapies for a wide range of conditions, such as various neurodegenerative disorders, cancers, cardiac, and autoimmune diseases (for reviews, see: Fleifel et al., 2018; Shroff et al., 2018; Solis et al., 2019; Song et al., 2018). Moreover, the reprogramming of somatic cells to induced pluripotent stem cells (iPSCs) (Okita et al., 2007; Takahashi et al., 2007) has revealed an enormous potential for the development of personalized cell therapies, as autologous stem cells may minimize immunological incompatibilities while circumventing the ethical dilemmas associated with using embryo-derived stem cells (ESCs).

Currently, a number of hurdles still hinder the development of safe and efficient therapies using PSCs (Martin, 2017). Despite the still unsolved issues, numerous cases of unproven and untested stem cell therapies involving adult stem cells as well as PSCs, frequently with harmful outcomes for the patients, have been reported (Bauer et al., 2018). A significant problem concerning the safety of PSC applications is teratoma formation from the transplanted cells, and various strategies have been tested to address this issue (Itakura et al., 2017; Nakamura and Okano, 2013). In this regard, the use of animal models to clarify potential hazards of PSC-based therapies in whole organisms is necessary. The common marmoset (*Callithrix jacchus*), a New World non-human primate, offers numerous advantages due to easiness of handling, absence of known zoonoses, high fertility, relatively short generation interval, and many close physiological similarities to humans. The development of age-related phenotypes similar to humans (Ross et al., 2012) makes the marmoset a suitable translational model for age-related conditions, such as Alzheimer’s (Tardif et al., 2011) and Parkinson’s disease (Yun et al., 2015). Gross morphometry heart characterization also identified the *Callithrix jacchus* as potential model for cardiac conditions (Senos et al., 2014). Development of efficient methods for genetic modifications in this species (Sasaki et al., 2009) has recently enabled the creation of several transgenic disease models for immunodeficiency, Parkinson’s, and polyglutamine diseases (Sato et al., 2016; Shimozawa et al., 2017; Tomioka et al., 2017).

Successful generation of marmoset iPSCs has been reported by a few research groups using retroviral (Tomioka et al., 2010; Wu et al., 2010) or transposon (Debowski et al., 2015) vectors. One disadvantage of these methods is the integration of the reprogramming transcription factors into the cell’s genome, which may cause genetic alterations. Reprogramming of marmoset cells to iPSCs has also been achieved with non-integrative episomal vectors (Vermilyea et al., 2017). Most recently, two research groups reported the derivation of marmoset iPSCs with synthetic mRNAs (Nakajima et al., 2019; Watanabe et al., 2019). Since mRNAs have short lifespan in the cells, multiple sequential transfections were necessary for successful reprogramming, subjecting the cells to increased stress. The reprogrammed iPSCs were maintained in undifferentiated state with MEF-conditioned medium (Watanabe et al., 2019) or on MEF feeders (Nakajima et al., 2019), underlining the need to optimize the xeno-free culture conditions for marmoset iPSCs, which is a prerequisite for the use of any therapeutic stem cells in preclinical trials.

In an alternative approach, the reprogramming transcription factors can be expressed by using self-replicating mRNA vectors based on the Venezuelan equine encephalitis virus (VEE-mRNAs). The 5’-capped, single-stranded viral mRNA contains four non-structural proteins followed by the structural viral proteins, which are transcribed from an internal S26 promoter and can be replaced with heterologous proteins of choice (Kinney et al., 1989; Petrakova et al., 2005). A significant advantage of this vector system is that a single transfection is sufficient for the long-term transgene expression in cells of different species. By construction of VEE-mRNA expression vector containing the human *OCT4, KLF4, SOX2*, and *c-MYC* (VEE-OKS-iM), one research group produced transgene-free human iPSCs (Yoshioka et al., 2013). To prevent mRNA elimination by the innate immune response, B18R recombinant protein (which binds and inhibits type I interferons) was used to maintain transgene expression until the completion of reprogramming.

In this report, we describe the generation of iPSCs from marmoset somatic cells by using self-replicating Tomato-modified VEE-OKS-iM mRNA. By selective inhibition with small molecule inhibitors, we first reprogrammed VEE-OKS-iM-iTomato-transfected marmoset fibroblasts to intermediate primary colonies with some neural progenitor characteristics, which we then converted to pluripotent stem cells by further culturing them in customized marmoset iPSC medium. These newly generated pluripotent cells are transgene-free and have been maintained long-term in feeder- and serum-free conditions. At the same time, they possess typical iPSC characteristics, such as expression of various pluripotency markers and the ability to differentiate into derivatives of the three primary germ layers *in vitro* and *in vivo*. Our study reveals a novel paradigm for flexible reprogramming of marmoset somatic cells that can be used to produce either cells of the neural stem cell lineage or iPSCs using the same primary colony population generated by a single transfection with VEE-mRNAs.

## RESULTS

### Derivation of primary colonies and iPSCs

The configuration of the modified T7-VEE-OKS-iM-iTomato plasmid and an outline of the reprogramming process are shown in Figures 1A and 1B. Successfully transfected marmoset fetal fibroblasts (cjFFs) were recognizable by their Tomato fluorescence (Tomato^+^) (Figure 1C). Following selection with Puromycin for 5-7 days, nearly all cells were Tomato^+^. Initially, we failed to obtain any iPSC lines when we cultured these cells on Geltrex-coated dishes with iPS-Brew or E8 medium, which prompted us to test different small molecule inhibitor supplements. In one of these experiments, primary colonies with compact morphology and clearly defined borders appeared in iPS-Brew supplemented with CHIR99021 and SB431542 (Figures 1D and 1E). These colonies were Tomato^+^ (Figure 1D), and varied widely in size between 200 and 3000 μm across. The primary colonies also exhibited alkaline phosphatase (AP) activity (Figure 1F). We manually picked 10-15 colonies/dish, disaggregated them in collagenase, and cultured them as single cells or small clumps in customized marmoset iPSC medium (iPS-Brew supplemented with IWR1, CHIR99021, CGP77675, rhLIF, and Forskolin). After 2-3 passages, we observed colonies with morphology characteristic of primate iPSCs (Figure 1G). These iPSC-like colonies were picked manually and further expanded and maintained by splitting with collagenase type IV (Figure 1H). In total, 4 iPSC lines (3 male and 1 female) were established from 3 different primary cjFF samples and have been maintained for 23 – 61 passages by the time of writing this report.

**Figure 1.**
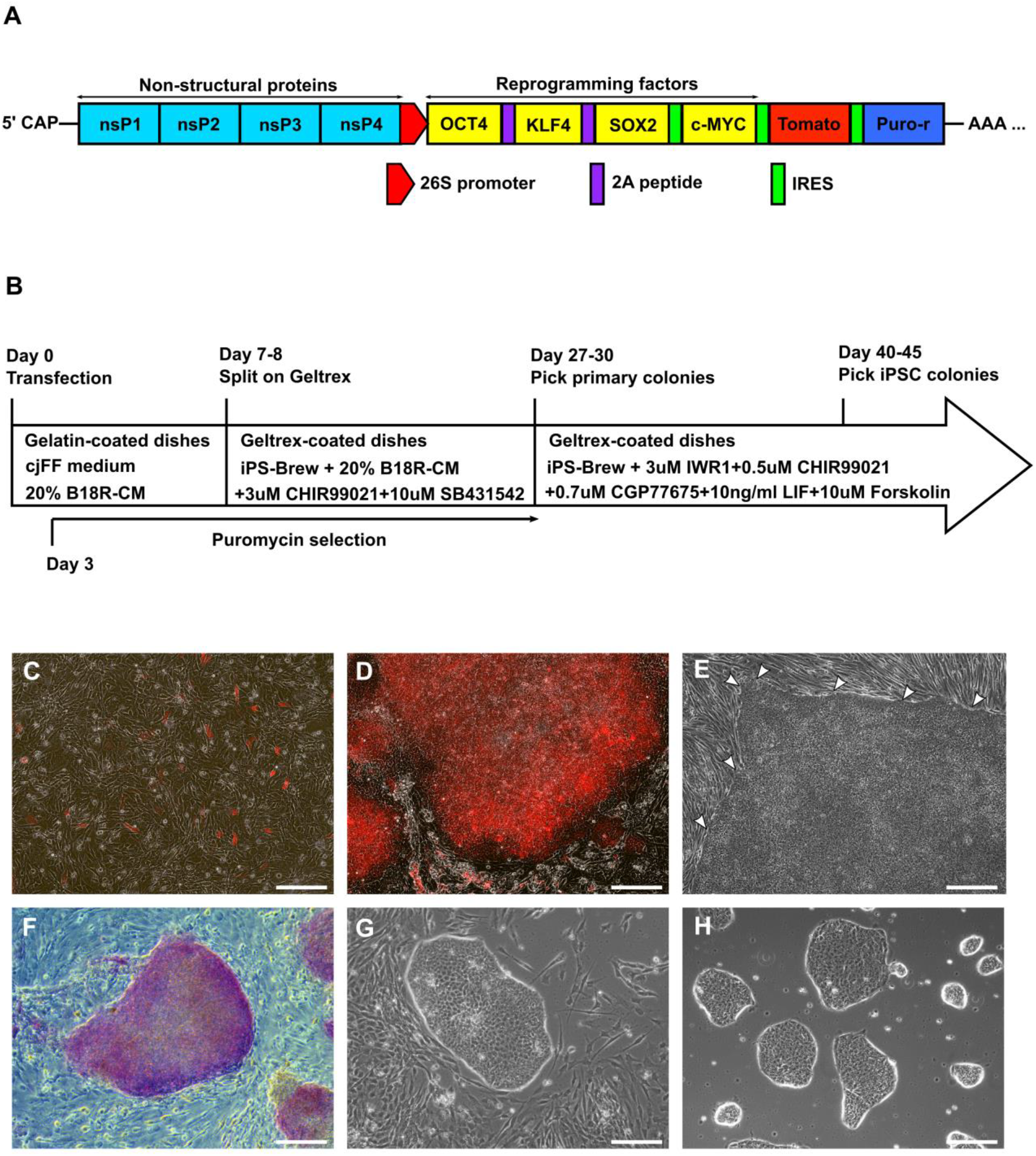
Generation of marmoset iPSCs with VEE-OKS-iM-iTomato. **A)** Structure of the reprogramming VEE-mRNA. **B)** Scheme of the reprogramming process. **C)** Image of cjFFs transfected with VEE-OKS-iM-iTomato at day 3 post-transfection. Transfected cells are recognizable by their red fluorescence. **D)** An intermediate primary colony at day 27 post-transfection with red VEE-OKS-iM-iTomato fluorescence. **E)** An intermediate primary colony at day 27 post-transfection with characteristic compact morphology and clearly defined borders (indicated with arrowheads). The cells beyond the upper and left border of the colony are non-reprogrammed cjFFs. **F)** An intermediate primary colony stained for AP. **G)** Marmoset iPSC colony after second round of reprogramming of the intermediate primary cells with IWR1, CHIR99021, CGP77675, LIF, and Forskolin. **H)** Morphology of marmoset iPSC colonies growing on Geltrex at P27. (All scale bars = 200 μm).

### Gene expression analysis

All iPSC colonies possessed strong AP activity (Figure S1A) and were shown to be positive for expression of OCT4A, NANOG, SSEA-4, TRA-1-60, TRA-1-81, CDH1, SALL4, and SOX2 by immunofluorescence (Figure 2A). Endogenous transcripts of *OCT4A* and *DPPA2* were detected by RT-PCR only in iPSCs, which clearly distinguished them from primary colonies and cjFFs (Figure 2B). Other pluripotency-related genes, such as *NANOG, GDF3, TERT, ZFP42, DPPA4*, and *CDH1* were detected at mRNA level in iPSCs as well as in primary colonies (Figure 2B). The expression of the exogenous reprogramming factors in intermediate primary colonies was confirmed at mRNA level by RT-PCR (Figure 2C). In absence of B18R-CM, all putative iPSC colonies were Tomator^−^ by P6, and the transgenes were not detected in iPSCs at mRNA or DNA level (Figure 2C). Although most pluripotency-related mRNAs were detected in primary colonies as well as iPSCs by conventional RT-PCR, real-time relative quantitation RT-PCR analysis revealed that *OCT4A, NANOG, GDF3, CDH1, LIN28*, and *DPPA4* were significantly upregulated in iPSCs relative to cjFFs and primary colonies (Figures 2D and 2E). Only *SALL4* expression in primary colonies was already in the range of *SALL4* expression in iPSCs (Figure 2E).

**Figure 2.**
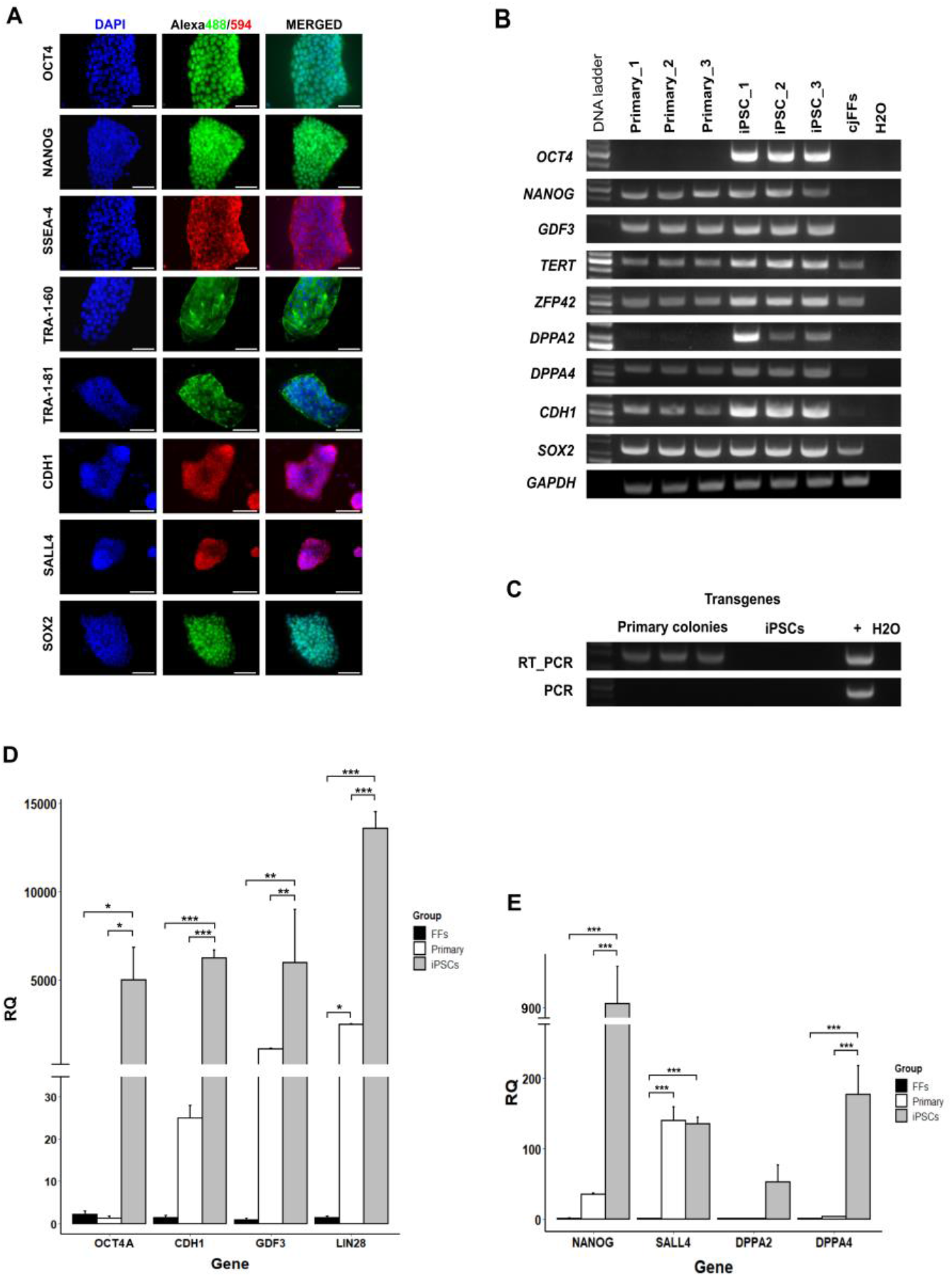
Expression of pluripotency-related genes in marmoset iPSCs. **A)** Immunofluorescence of iPSC colonies stained for expression of OCT4, NANOG, SSEA-4, TRA-1-60, TRA-1-81, CDH1, SALL4, and SOX2. (Scale bars = 50 μm). **B)** RT-PCR analysis of endogenous pluripotency marker expression in primary cultures and iPSCs. **C)** RT-PCR and PCR detection of transgenes in primary colonies and iPSCs. Plasmid DNA used as positive control (denoted as “+”). **D)** Real-time RT-qPCR analysis of relative expression levels of *OCT4A*, *CDH1*, *GDF3*, and *LIN28* in cjFFs, intermediate primary colonies, and iPSCs. **E)** Comparison of the relative expression levels of *NANOG*, *SALL4*, *DPPA2*, and *DPPA4* in cjFFs, intermediate primary colonies, and iPSCs (each group n = 3) by real-time RT-qPCR (RQ = 2^−ddCt^). (Data are presented as mean + SEM. Statistically significant differences between different experiment groups are indicated by asterisks as follows: * p < 0.05; ** p < 0.01; *** p < 0.001.)

Neural markers *NESTIN, NOTCH1*, *GBX2* as well as neural plate border markers *PAX3* and *MSX1* were upregulated in intermediate primary colonies relative to cjFFs (Figure 3A, 3B, and 3C). Following the second reprogramming step, the neural markers *NESTIN*, *NOTCH1*, *CDH2*, *PAX3*, *MSX1*, and *DLX5* became significantly downregulated in the iPSCs (Figures 3B and 3C). At the same time, the iPSCs significantly upregulated *SOX2* and *OTX2* relative to both cjFFs and primary colonies (p-values < 0.001 and < 0.03, respectively) (Figure 3A). The expression of OTX2 in iPSCs was also confirmed by immunofluorescence (Figure S1B). Finally, genes associated with endo- or mesodermal differentiation *GATA6, MIXL1*, and *NKX2.5*, were significantly downregulated in primary colonies and iPSCs relative to cjFFs (Figure 3D) (all p-values ≤ 0.01). Other differentiation genes, such as *HAND1*, *GATA4*, and *TBX5* also showed downregulation in primary colonies and iPSCs (Figure 3D); however, it was not found statistically significant. Altogether, the analysis of the pluripotency-related and lineage-specific markers allows an accurate discrimination between the fetal fibroblasts, the primary intermediate colonies, and the putative iPSCs. Furthermore, the neural marker analysis indicates an ectodermal/ neuronal identity of the primary intermediate colonies.

**Figure 3.**
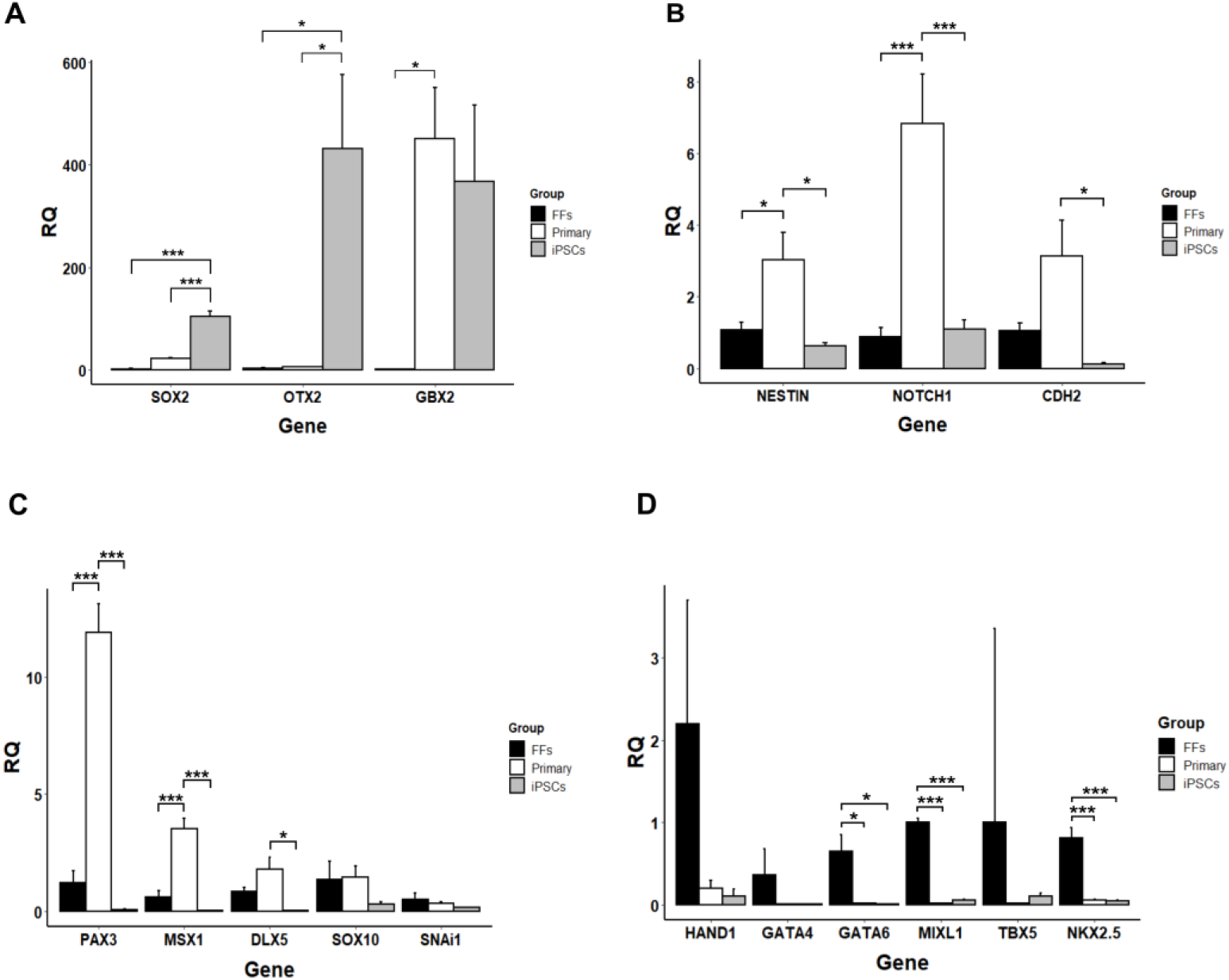
Real-time RT-qPCR analysis of lineage-specific marker expression. **A)** Relative expression of *SOX2*, *OTX2*, and *GBX2*. **B)** Relative expression of neural stem cell markers *NESTIN*, *NOTCH1*, and *CDH2* in cjFFs, intermediate primary colonies, and iPSCs by real-time RT-qPCR. **C)** Relative expression of neural crest markers *PAX3, MSX1, DLX5, SOX10*, and *SNAi1*. **D)** Relative expression of endo/mesodermal markers *HAND1*, *GATA4, GATA6, MIXL1, TBX5*, and *NKX2-5* (RQ = 2^−ddCt^). (Data are presented as mean + SEM. Each biological group n = 3. Significant differences are indicated with asterisks as follows: * p < 0.05; ** p < 0.01; *** p < 0.001.)

### In vitro and in vivo differentiation of marmoset iPSCs

When cultured in suspension, the iPSCs formed EBs, some of which became cavitated within 7 days (Figure S1C). A small number (2-3%) of the EBs also showed contracting activity due to presence of immature cardiomyocytes (Video S1). On Geltrex-coated glass coverslips the EBs formed outgrowths containing β-III-Tubulin-positive neurons (Figure S1D) and SOX17-positive endoderm-like cells (Figure S1E). We performed directed neural differentiation by first “neuralizing” the iPSCs with dual SMAD inhibition (dorsomorphin and SB431542) followed by splitting and culture in NSC medium, where the cells maintained expression of NSC markers PAX6 and Nestin (Figure S1F.). When these NSC-like cells were cultured in neurobasal medium on poly-ornithine and laminin, they formed neural rosettes (Figure S1G) and subsequently differentiated into neurons (Figure S1H) positive for β-III-Tubulin and NESTIN (Figures 4A and 4B). Directed differentiation of the iPSCs into endoderm resulted in appearance of typical “cobblestone”-like cell layer (Figure 4C), where approximately 40% of the cells stained positive for AFP at different intensities by immunofluorescence (Figure 4D). Finally, directed cardiac differentiation resulted in contracting myocytes organized in several clusters (Video S2). These primitive cardiomyocytes were metabolically selected, transferred on glass coverslips, and stained for expression of cardiac-specific proteins. The majority of the cardiomyocytes were positive for cTnT, MLC2a, α-actinin, CX43, and Titin (Figures 4E-H).

**Figure 4.**
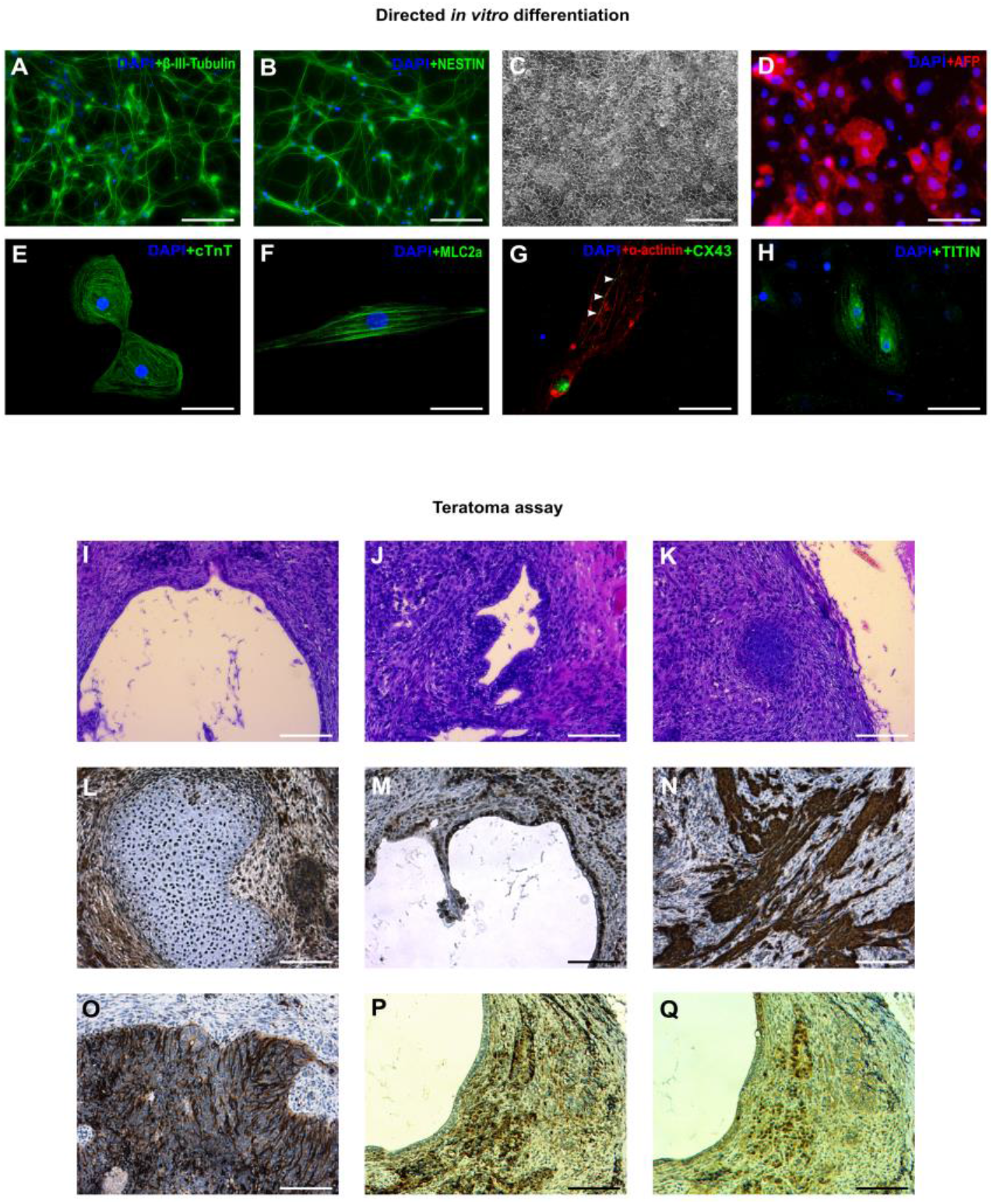
In vitro and in vivo differentiation of marmoset iPSCs. **A** and **B)** Neurons positive for β-III-Tubulin and NESTIN. **C)** Endodermal differentiation with cobblestone-like morphology. **D)** Primitive endoderm stained with anti-AFP. **E-H)** Cardiomyocytes stained for: **E)** cTnT, **F)** MLC2a, **G)** α-actinin and CX43 (arrowheads), and **H)** Titin. **I-Q) Teratoma assay: I)** Gut-like endoderm. **J)** Gland-like endoderm. **K)** Bone. **L)** Bone tissue stained for SOX17 (which is also involved in osteogenesis). **M)** Gut-like tissues stained with anti-SOX9. **N)** Smooth muscle stained for SMA. **O)** Neuronal-like cells stained for beta-III-Tubulin. **P** and **Q)** Successive sections stained for presence of NESTIN and PAX6. (Scale bars: A, B, D, E, F, G, H = 50 um; C = 200 μm; I-Q = 100 μm).

When injected into immunodeficient mice, the iPSCs formed teratomas with presence of primitive gut-like endoderm (Figure 4I), gland-like structures (Figure 4J), and bone tissues (Figures 4K and 4L). The endodermal cells were positive for SOX9 (Figure 4M), while presence of mesoderm was confirmed by SMA staining (Figure 4N). Ectodermal differentiation was demonstrated by presence of β-III-Tubulin, PAX6, and NESTIN-positive cells (Figures 4O-Q). In summary, these data demonstrate pluripotency of the iPSCs.

### Karyotyping

Three iPSC lines were processed for karyotyping and at least 25 images/ line where all chromosomes were clearly distinguishable were used for counting. All lines had normal chromosome numbers where 2 were male (46 XY), and one was female (46 XX). (Figures S1I – S1K).

### Neurogenic potential of the intermediate primary colonies

When the intermediate primary colonies were mechanically picked, broken to small fragments, and cultured in NSC medium, multiple neural rosette-like structures formed in areas with high cell density (Figure 5A). These rosettes were picked, disaggregated, and cultured as monolayer in NSC medium (Figure 5B). For the entire period of culture (7-10 passages) the rosette-derived cells maintained SOX2 and NESTIN expression (Figure 5C). Differentiation was conducted by first generating neurosphere-like aggregates in suspension for 7 days (Figure 5D) and then culture on poly-ornithine and laminin, whereupon the cells differentiated into neurons (Figure 5E) expressing β-III-Tubulin and MAP2 (Figure 5F). We conclude from these data that the intermediate colonies have neurogenic potential when cultured under appropriate conditions.

**Figure 5.**
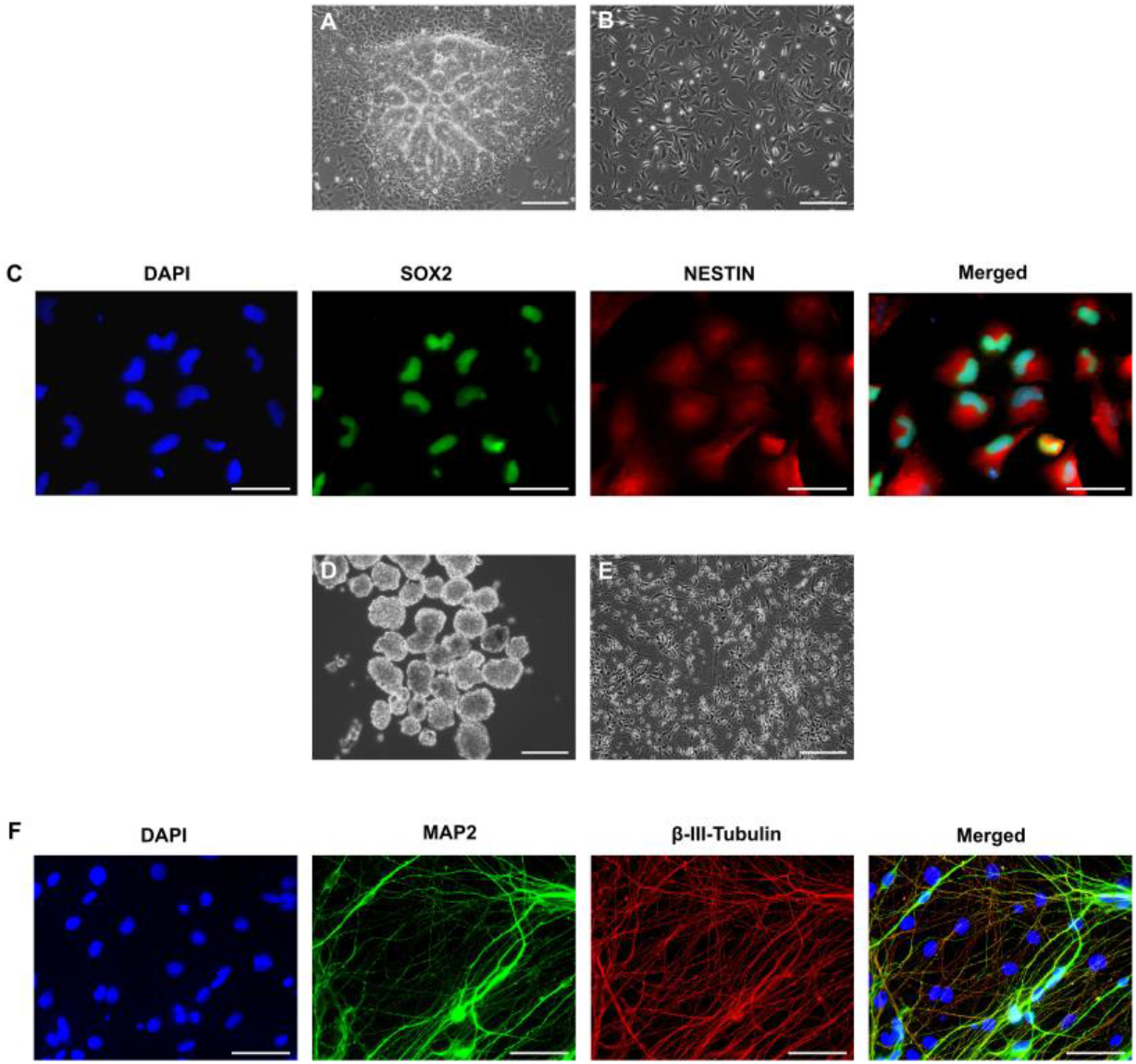
Directing the intermediate primary colonies into the neural lineage. **A)** Neural rosettes in P1 after culture in NSC medium containing bFGF and EGF. NSC-like cells growing on Geltrex in NSC medium. **C)** Neural sphere-like aggregates in suspension culture. **D)** Neuronal cells at the end of differentiation. **E)** NSC-like cells expressing SOX2 and NESTIN. **F)** Immunofluorescence from neurons stained with anti-MAP2 and β-III-Tubulin. (Scale bars: A-D = 200 μm; E and F = 50 μm).

## DISCUSSION

Among the non-integrative methods for iPSC production, episomal plasmids have been a popular choice due to their relatively lower costs and simplicity of use. One of the few caveats of this expression system is that the episomes currently used by most groups contain immortalization factors, such as large T antigen (Yu et al., 2009) or shRNA for p53 knockdown (Okita et al., 2011), which in some cases have been shown to cause genomic instability (Schlaeger et al., 2015; Tidball et al., 2016). In addition, episomal plasmid expression may persist in up to 33% of the reprogrammed cells even after P11 (Okita et al., 2011; Schlaeger et al., 2015). Using mRNAs for iPSC generation eliminates the risks of genomic integration and allows for quick and efficient removal of the transgenes. At the same time, the short half-life of the mRNA messages in the cells is a major disadvantage, since a sustained expression of the reprogramming factors over time is necessary for successful reprogramming. For this reason, non-replicating mRNAs need to be introduced multiple times into the cells. In order to mitigate the low reprogramming efficiency, additional reprogramming factors have been included: various miRNAs (Nakajima et al., 2019), or *NANOG* and *LIN28* transgenes as well as shRNA-p53 (Watanabe et al., 2019). In contrast to these reports, we were able to derive marmoset iPSCs with a single transfection of self-replicating VEE-mRNAs carrying only the four Yamanaka factors (*OCT4, KLF4, SOX2*, and *c-MYC*) and, importantly, devoid of any immortalization factors. However, it must be noted that this was possible only after modification of the protocol to include an initial reprogramming step with generation of partially reprogrammed intermediate primary colonies by culture with GSK3β and ALK5 inhibitors (CHIR99021 and SB431542, respectively), because protocols optimized for reprogramming of human cells failed to produce any marmoset iPSCs. Under the influence of the two small molecule inhibitors, the transfected cjFFs initiated robust proliferation and formed multiple primary colonies, which we at first mistook for iPSCs due to their compact morphology and clearly defined borders. Subsequently, we found out that these cells were negative for endogenous OCT4A but expressed some neural progenitor markers (SOX2, NESTIN, NOTCH1, CDH2, GBX2) as well as early neural plate border specifiers (PAX3 and MSX1). These findings prompted us to direct these cells into the neural lineage and we were successful in producing SOX2^+^ and NESTIN^+^ cells that could be maintained in NSC culture conditions and were capable of neuronal differentiation. The further characterization of these cells is currently ongoing and the results will be published in a separate report.

When cultured further with modified iPSC medium containing Wnt signaling inhibitor IWR1, low concentration of TGF3β inhibitor CHIR99021, and Src kinase inhibitor CGP77675, the intermediate primary colonies formed iPSC colonies within 2-3 passages. This conversion was dependent on the presence of IWR1, which has been shown to stabilize AXIN1/2 and cause cytoplasmic retention of β-catenin by preventing its translocation to the nucleus (Jho et al., 2002). It has been demonstrated that cytoplasmic β-catenin interacts with TAZ to promote self-renewal of mouse epiblast stem cells as well as human iPSCs (Zhou et al., 2017). In addition, together with CHIR99021, Src kinase inhibitor CGP77675 has been shown to maintain the developmental competence of mouse ESCs (Choi et al., 2017). We found that CGP77675 was important for the generation of marmoset iPSCs under feeder-free conditions as well as for their long-term maintenance, as its withdrawal from the culture medium later caused differentiation (results not shown). Lastly, although rhLIF and Forskolin were not necessary for the derivation of marmoset iPSCs, they bolstered the reprogramming process by improving cell proliferation. Under the influence of the inhibitor cocktail, many endogenous pluripotency markers became upregulated, including OCT4A, which was not detectable in cjFFs or primary colonies. At the same time, neural progenitor markers were downregulated in the iPSCs, with the exception of SOX2 and OTX2. Both markers play important roles in the development of the nervous system; however, SOX2 is also involved in pluripotency, while OTX2 is expressed in the epiblast and is characteristic of mouse epiblast stem cells and human ESCs (Acampora et al., 2013). Therefore, the relatively high expression of OTX2 in our marmoset iPSCs suggests that they are primed pluripotent cells.

The reprogramming protocol employed in this study differs from the majority of protocols described to date for the production of iPSCs from NHPs and other species. One advantage of this novel approach is the generation in the first reprogramming step of multiple primary colonies, which then can be flexibly used for the derivation of neural and/or pluripotent cells, according to the objectives of the particular study (Figure 6). Despite providing a proof-of-principle, it remains to be seen whether this paradigm would also function in cells of species other than the common marmoset. Nevertheless, taking into consideration the data obtained from this study, we expect that our results will help to further improve the conditions for reprogramming and culture of marmoset iPSCs. We also expect that our new reprogramming method, resulting in transgene-free iPSCs, in combination with the fully defined culture medium, will enhance the use of the marmoset monkey as a NHP species in translational studies of regenerative medicine. Finally, the relatively efficient genetic modification of marmoset monkeys allows the testing of key findings made in stem cell culture in long-term *in vitro* embryo culture (Ma et al. 2019; Niu et al. 2019) or even *in vivo*.

**Figure 6.**
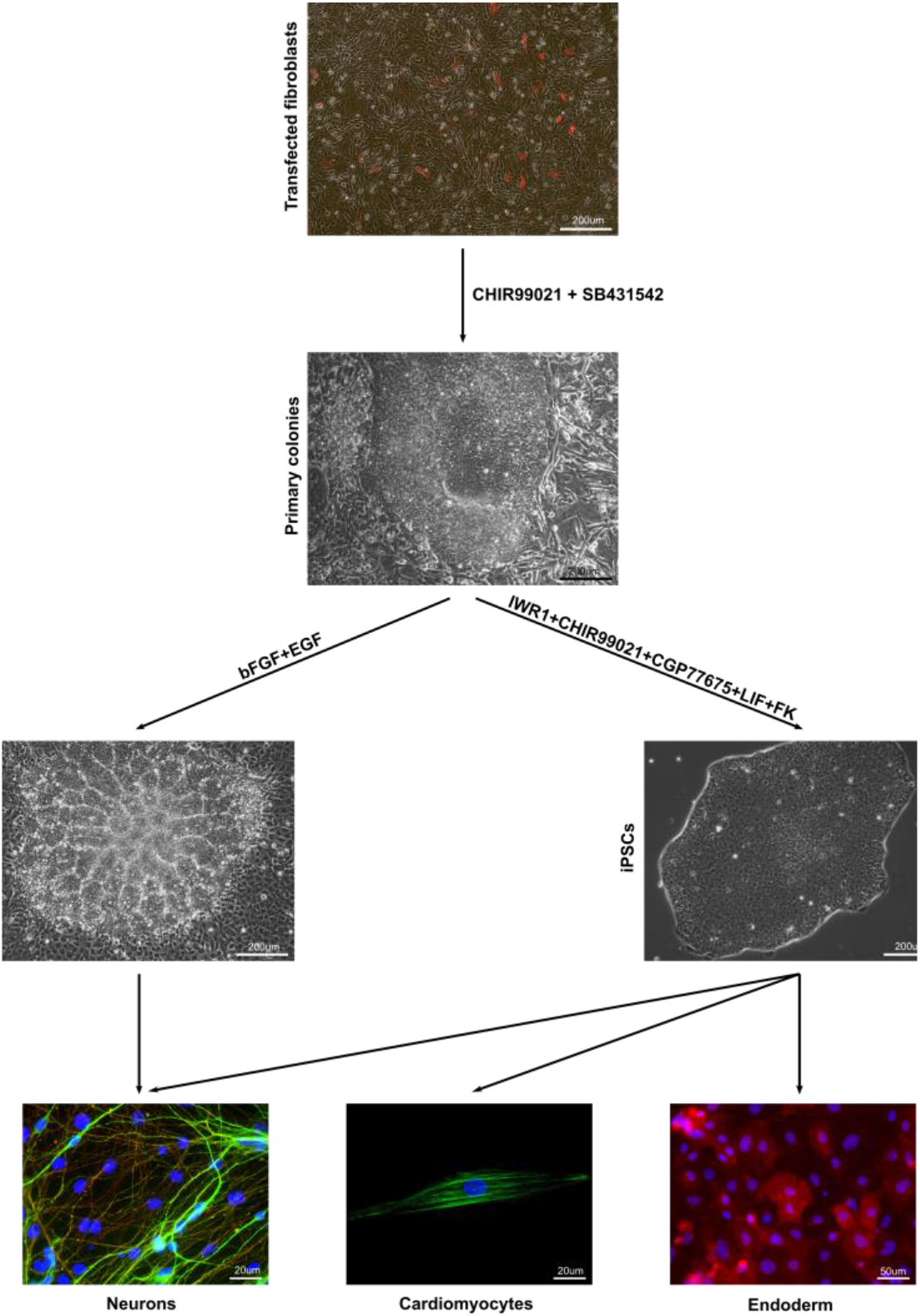
Paradigm for two-step flexible reprogramming of marmoset somatic cells to pluripotency or to the neural lineage. Transfection of somatic cells with vector carrying reprogramming transcription factors and culture in medium containing CHIR99021 and SB431542 leads to generation of intermediate primary colonies. These colonies can be directed into the neural lineage by culture with bFGF and EGF, or to be further reprogrammed to iPSCs by culture with IWR1, CGP77675, LIF and Forskolin.

## CONCLUSIONS

By using self-replicating VEE-mRNAs and small molecule inhibitors, we successfully derived transgene-free marmoset iPSCs in feeder- and serum-free conditions. By inhibition of GSK3β and the TGF3β receptor signaling we first established intermediate primary colonies with capability to differentiate into the neural lineage. Subsequently, we converted these primary colonies to iPSCs by modulating the β-catenin signaling and by inhibiting the Src kinase. With our experiments, we established a novel paradigm for flexible reprogramming of marmoset somatic cells that will enhance the role of the common marmoset as biomedical non-human primate model for the development of stem cell therapies.

## EXPERIMENTAL PROCEDURES

### Isolation of marmoset fetal fibroblasts (cjFFs)

Animal care as well as all treatment procedures were in accordance with the current regulations as outlined in the animal protection law and reflected in the institutional guidelines. Marmoset cjFFs were isolated from leftover fragments of marmoset fetuses from days 70-74 of gestation, first used in other unrelated projects that have been fully approved by the Lower Saxony’s State Office of Consumer Protection and Food Safety (LAVES) (license numbers 42502-04-16/2129 and 42502-04-16/2130) including a positive ethics evaluation. Tissues of the fetal dorsal body wall and limb buds were finely minced with scalpel blade and incubated in a mixture of 1:1 (v:v) Accutase (Gibco) and Collagenase IV (Worthington Biochemical Corporation)(2 mg/ml) at 37°C for 15 min, followed by trituration to disaggregate the tissues to single cells or small clumps. The suspension was centrifuged at 300 g for 5 min, re-suspended in M20 culture medium (Dulbecco’s DMEM containing GlutaMAX (Gibco) supplemented with 20% FBS (Gibco), non-essential amino acids (NEAA (Gibco), penicillin/streptomycin (Gibco), and 5 ng/ml bFGF), and plated on gelatin-coated 10 cm tissue culture dishes. The cells were subsequently split 1:4-1:6 with Accutase every 3-4 days and maintained in M20 medium.

### Synthesis of self-replicating mRNAs (VEE-OKS-iM-iTomato)

The plasmid T7-VEE-OKS-iM was a gift from Dr. Steven Dowdy (Addgene plasmid # 58972 ; http://n2t.net/addgene:58972 ; RRID:Addgene_58972) and has been previously used by his research group for the generation of human iPSC (Yoshioka et al., 2013). Using conventional molecular cloning techniques, an IRES-Tomato dsDNA fragment (iTomato) was inserted into the *NotI* restriction site between the stop codon of c-MYC and the IRES site of the puromycin resistance gene. The resulting plasmid (T7-VEE-OKS-iM-iTomato) was linearized by MluI digest and mRNA was *in vitro* transcribed, capped, and polyadenylated using HiScribe ARCA T7 mRNA Kit (with tailing) (New England Biolabs) according to the manufacturer’s instructions.

### Production of B18R-conditioned medium (B18R-CM)

The B18R-6His construct was excised from plasmid pTNT-B18R-6His (also gift from Dr. Steven Dowdy (Addgene plasmid # 58979; http://n2t.net/addgene:58979; RRID:Addgene_58979)) and ligated into a *PiggyBac* transposon plasmid pTT-PB-Puro^r^, (Debowski et al., 2015) by using standard molecular biology techniques, to generate pTT-PB-B18R-6His-Puro^r^ for genomic integration and constitutive protein expression. For production of B18R-CM, 2 ×10^6^ mitotically active mouse embryonic fibroblasts (MEFs) were co-transfected with 2 μg pTT-PB-B18R-6His-Puro^r^ and 1 μg pcA3-PBase-Tomato using Lipofectamine 3000 per manufacturer’s instructions. Following selection with 1 μg/ml puromycin (Sigma-Aldrich) for 2 weeks, the B18R-transgenic MEFs were split at 1×10^6^ cells/dish in 10 cm culture dishes and grown with M10 medium until reaching 100% confluency, and then for another 2 days following change of medium. The first and second conditioned media were collected, mixed, centrifuged at 4000 g for 10 min to pellet cells and debris, filtered through 0.4 μm filters, and stored at −80° C until use. The B18R-CM was added at 20% (v:v) to the cjFF culture medium 2-3 days before transfection with mRNAs as well as to the reprogramming medium thereafter.

### Electroporation of cjFFs with VEE-OKS-iM-iTomato and generation of intermediate primary colonies

The cjFFs were transfected at P4-6 with Multiporator (Eppendorf) according to the protocol provided by the manufacturer. Briefly, cFFs were suspended in hypoosmolar electroporation buffer (Eppendorf) at a concentration of 2×10^6^ cells/ml and incubated at room temperature for 20 min. VEE-OKS-iM-iTomato mRNA was added at 6 μg/380 μl suspension, the cells were transferred to electroporation cuvette with 2 mm gap (Sigma-Aldrich) and electroporated at 550 V, 100 μs, 1 square pulse. Following 10 min recovery at room temperature, the cells were transferred to 3 gelatin-coated wells (6-well plate format) and cultured at 37°C with M20 medium supplemented with 20% B18R-CM for 9-10 days. To select the cells expressing the reprogramming mRNA, 1 μg/ml Puromycin (Sigma-Aldrich) was added starting at 48-72 hours post-transfection (p.t.) and was maintained till appearance of the intermediate primary colonies. After 7-8 days the selected cells were seeded on Geltrex-coated 6 cm tissue culture dishes (CytoOne) at 20-25×10^5^ cells/dish and cultured in primary reprogramming medium (StemMACS iPS-Brew XF medium (iPS-Brew; Miltenyi Biotech) supplemented with 3 μM CHIR99021 (Sigma-Aldrich) and 10 μM SB431542 (Selleckchem) and 20% B18R-CM until the picking of the intermediate primary colonies (27-30 days p.t.).

### Generation and culture of marmoset iPSCs

Intermediate primary colonies with compact morphology were manually picked, digested in 1mg/ml collagenase type IV solution (Worthington Biochemical Corporation) and disaggregated by pipetting to single cells or small clumps. The cell suspension was seeded in 6 cm Geltrex dishes (30-40×10^5^ cells/dish) in marmoset iPSC medium (iPS-Brew supplemented with 3 μM IWR1, 0.5 μM CHIR99021, 0.7 μM CGP77675, 10 ng/ml hrLIF, and 7 μM Forskolin) and cultured for 2-3 passages in hypoxic conditions (5% O_2_, 5% CO_2_, and 90% N_2_) until the appearance of colonies with iPSC-like morphology. These colonies were manually picked, disaggregated in collagenase solution to small clumps and passed to fresh Geltrex dishes in iPSC medium with CGP77675 concentration reduced to 0.3 μM. The iPSCs were further maintained by splitting with collagenase at ratio 1:3-1:4 every 3-4 days. To increase survival, 5 μM Pro-survival Compound (Merck Millipore) was added only on the day of splitting. Hypoxic conditions were maintained during the entire duration of iPSC culture.

### Directing intermediate primary colonies into the neural lineage

Primary colonies were picked manually, broken to fragments (50-100 μm) and plated on Geltrex-coated dishes in NSC medium consisting of neurobasal medium (DMEM/F12 (ThermoFisher) supplemented with L-glutamine, 1% N2, 2% B27, and 50 μg/ml L-ascorbic acid (Sigma-Aldrich)) supplemented with 20 ng/ml bFGF (Peprotech), and 20 ng/ml EGF (Peprotech). The neural rosette-like areas were manually picked, disaggregated in Accutase, and cultured further in the same conditions. For neural differentiation, NSC-like cells at 100% confluency were scraped with 100μl pipette tip and the cell clumps were cultured in suspension in 60 mm Petri dishes in neurobasal medium without bFGF and EGF for 7 days. The resulting neurosperes were cultured on poly-L-ornithine and laminin-coated dishes for 7 days and then split with Accutase on glass coverslips for immunofluorescence.

### Total RNA isolation, PCR, RT-PCR, and real-time relative quantitation

Total RNA was purified using Nucleospin RNA Plus kit (Macherey-Nagel) according to the manufacturer’s protocol. The extracted RNA was treated with DNAseI (New England Biolabs) and reverse transcription was performed with Omniscript RT Kit (Qiagen). The PCR amplifications were conducted using Taq DNA Polymerase with Standard Taq buffer (New England Biolabs). The nucleotide sequences for RT-PCR and real-time PCR primers (synthesized by Sigma-Aldrich) are shown in Table 1. Relative quantitation real-time qPCR was performed with StepOnePlus Real-time PCR system (Applied Biosystems) using Power SYBR Green PCR master mix (Applied Biosystems). Three independent biological samples from each experimental group (cjFFs, Primary, and iPSCs) with at least two technical replicates/ sample were included in each analysis. Statistical analysis (one-way ANOVA with Tukey HSD test) and graphical presentation of the results were performed with R-studio.

**Table 1.**
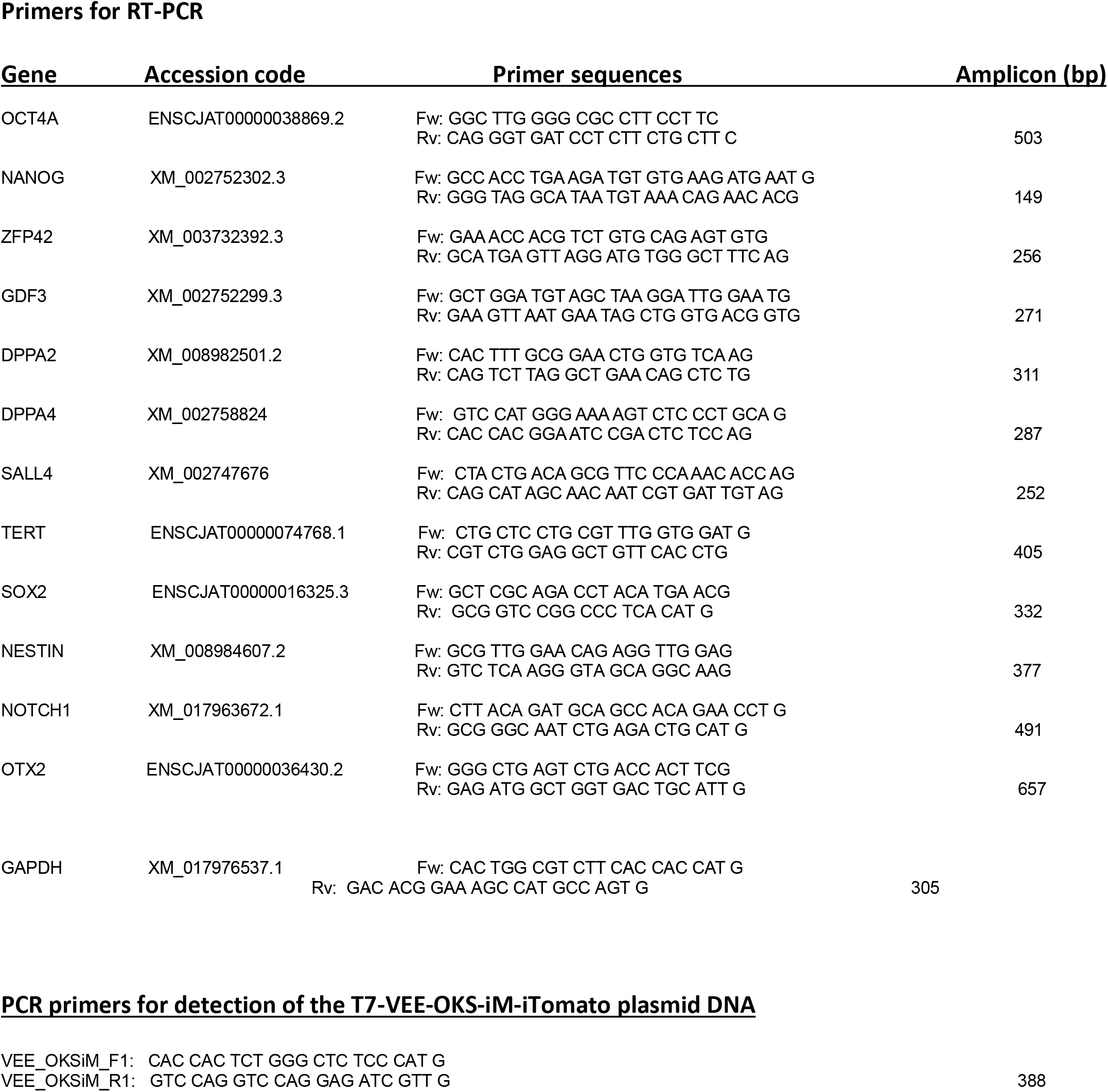
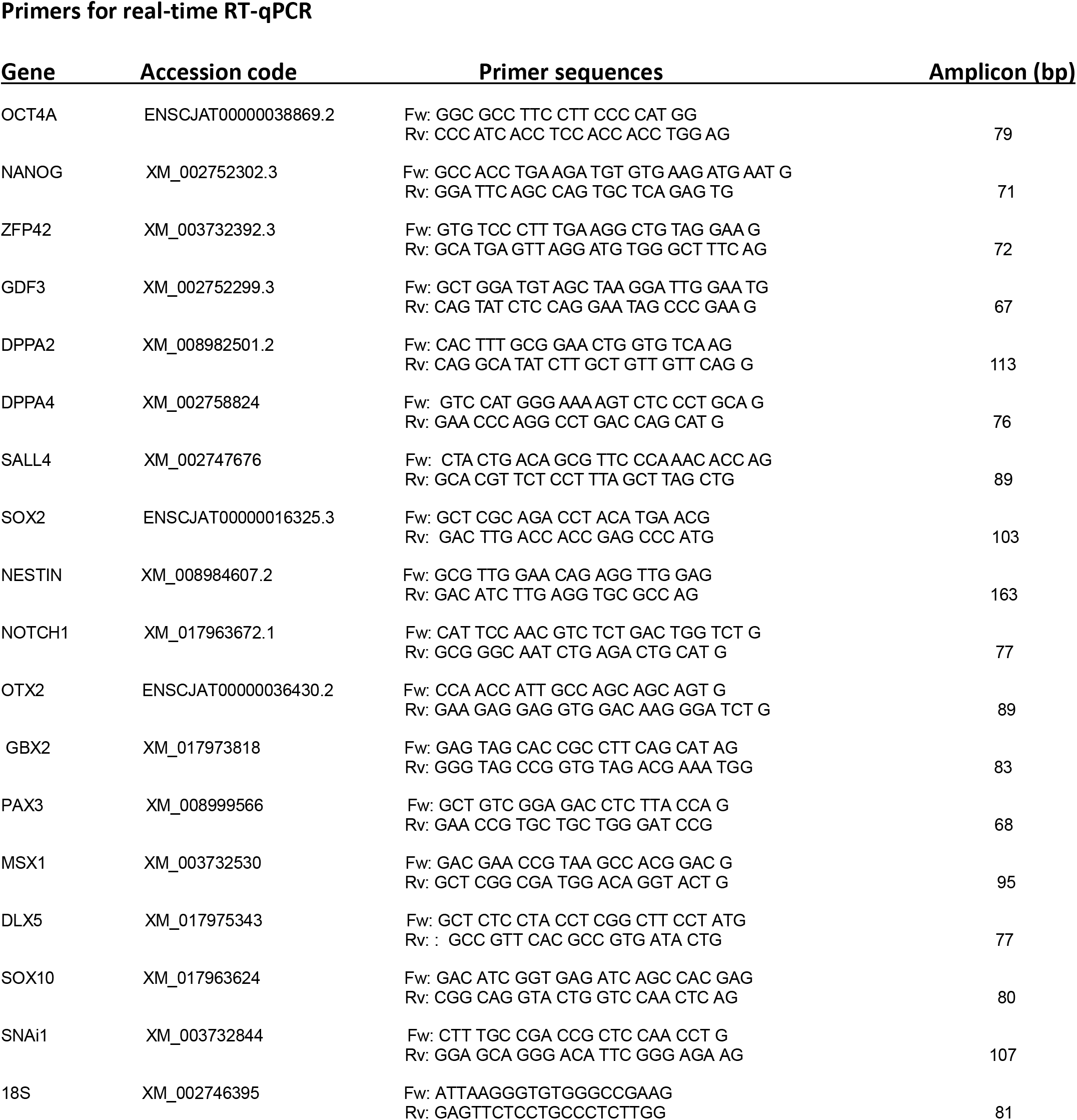
Primer sequences.

### Immunofluorescence and alkaline phosphatase activity

All procedures were performed at room temperature. The iPSC intended for immunofluorescence analysis were cultured on Geltrex-coated glass coverslips. The cells were fixed in 4% paraformaldehyde solution for 15 min., permeabilized with 0.15% Triton X for 10 min., and blocked with 1% BSA for 1 hour prior to incubation with primary antibodies raised against OCT4 (Cell Signaling, Cat. # 28901:800), NANOG (Cell Signaling, Cat. # 4903; 1:500), SOX2 (Cell Signaling, Cat. # 3728; 1:200), SSEA-4 (Millipore, Cat. # MAB4304; 1:100), TRA-1-60 (eBioscience, Cat. # 14-8863; 1:100), TRA-1-81 (eBioscience, Cat. # 14-8883; 1:100), SALL4 (Sigma-Aldrich, Cat. # HPA015791; 1:200), CDH1 (Cell Signaling, Cat. # 3195; 1:400), or OTX2 (Sigma-Aldrich, HPA000633; 1:200). Differentiated endodermal or neuronal cells were processed as described above and immunostained with anti-β-III-Tubulin (Sigma-Aldrich, Cat. # T8660; 1:100), anti-NESTIN (MerckMillipore, Cat. # MAB5326; 1:100), anti-MAP2 (Sigma-Aldrich, Cat. # HPA012828; 1:100), anti-SOX17 (ThermoFisher, Cat. # PA5-23352; 1:200), or anti-alpha Fetoprotein (AFP) (DAKO, Cat. # A0008; 1:300). Cardiomyocytes were immunostained with anti-α-Actinin (Sigma-Aldrich, Cat. # A7811; 1:100), anti-cTnT (ThermoFisher, Cat. # MS-295-PABX; 1:200), anti-MLC2a (Synaptic Systems, Cat. # 311-011; 1:200), anti-Titin (MerckMillipore, Cat. # MAB1553; 1:50), and anti-CX43 (Abcam, Cat. # ab11370; 1:1000). Visualization was performed with Alexa Fluor 488 donkey anti-rabbit, Alexa Fluor 488 goat anti-mouse, or Alexa Fluor 594 donkey anti-mouse secondary antibody (Invitrogen) (All diluted 1:1000). Alkaline phosphatase activity was determined using Alkaline Phosphatase kit (Sigma-Aldrich) as instructed by the manufacturer.

### In vitro differentiation of marmoset iPSCs

Embryoid bodies were generated by treating iPSC cultures with Versene (Gibco) for 3 min at room temperature, scraping off and culturing the cell clumps in Petri dishes in Iscove’s medium (IMDM with GlutaMAX (ThermoFisher) supplemented with 20% FBS, NEAA, and 450 μM 1-monothyoglycerol (Sigma-Aldrich)) for 7-10 days. The EBs were then cultured on Geltrex-coated glass coverslips for another week and processed for immunofluorescence. Directed cardiomyocyte differentiation was performed with iPSCs at P9-12 as described by Tiburcy et al., 2017. Neural differentiation was achieved by neuralization in DMEM/F12 supplemented with 10% Knockout Serum Replacement (ThermoFisher), NEAA, 50 μg/ml ascorbic acid (Sigma-Aldrich), 2 μM SB431542 (Selleckchem), and 1.5 μM dorsomorphin (Selleckchem) for 5-7 days. Then, the cells were split with Accutase and cultured in NSC medium. To differentiate into neurons, the cells were cultured on poly-L-ornithine and laminin-coated dishes or glass coverslips in neurobasal medium for 10-14 days. For differentiation into endoderm, the iPSCs were first cultured in basal medium (RPMI-1640 with 2% B27) supplemented with 100 ng/ml Activin A (Peprotech) and 3 μM CHIR99021 for 3 days and then with 5 ng/ml bFGF, 20 ng/ml BMP4 (Peprotech) and 0.5% DMSO (Sigma-Aldrich) for 5 more days. At the end of experiment the differentiated cells were passed on Geltrex-coated glass coverslips with Accutase and immunostained with anti-AFP antibodies.

### Teratoma assay

Marmoset iPSCs at P20-30 were harvested with collagenase and re-suspended in cold Geltrex solution prepared with iPSC culture medium at concentration of 1×10^7^ cells/ml together with 1×10^5^/ml mitotically-inactivated mouse fetal fibroblasts (MEFs). The cells were transported on ice to the mouse housing facility where they were injected subcutaneously into the left flank of immunodeficient SCID/beige mice (C.B-17/IcrHsd-scid-bg) at 100μl/injection/mouse. The mice were bred in the central facility for animal experimentation at the University Medical Center Göttingen under specific pathogen-free conditions in individually ventilated cages and in a 12 h light-dark cycle. The mouse experiments had been approved by the local government (33.9-42502-04-19/3074) and were carried out in compliance with EU legislation (Directive 2010/63/EU). Mice with tumor growth (detected by regular palpation) were sacrificed before the tumors reached 1 cm size approximately 1-3 months after injection. All remaining animals were sacrificed after three months and autopsies were performed to exclude an unrecognized tumor growth.

The paraffin embedding, sectioning, and immunohistochemical staining of the teratomas were performed as described previously (Debowski et al., 2015). The presence of lineage-specific markers was determined by incubation with anti-β-III-Tubulin, anti-NESTIN, anti-PAX6 (MerckMillipore, Cat. # AB2237; 1:500), anti-smooth muscle actin (SMA) (Sigma-Aldrich, Cat. # A2547, 1:1000), anti-SOX17, and anti-SOX9 (MerckMillipore, AB5535, 1:300) antibodies and visualization with EnVision FLEX Mini kit (DAKO).

### Karyotyping

The chromosomal numbers of four iPSC lines were determined at P21-50 as described earlier (Petkov et al., 2018). Briefly, the cultures were arrested at metaphase by adding 0.02 mg/ml Demecolcine (Sigma) to the culture medium for 1.5 hours and then disaggregated with Accutase to single cells. Following incubation in hypoosmotic 0.56% KCl solution for 20 min., the cells were fixed in 3:1 methanol/acetic acid (v:v) fixative and metaphase spreads were generated by dripping 50 μl drops of the cell suspension on glass slides positioned at a slight angle over a steaming water bath. The metaphase spreads were air-dried, stained in Giemsa solution (Sigma) for 10 min., and photographed with Axiophot microscope (Zeiss) equipped with CRI Nuance multispectral imaging camera. Approximately 25-30 images per sample were used for chromosome counting.

## AUTHOR CONTRIBUTIONS

Experimental design: SP and RB. Performed the reprogramming experiments and characterization of the cells: SP. Teratoma assays: RD. Directed cardiac differentiation and analysis: IR-P. Manuscript writing: SP, RD, IR-P, and RB.

## ACKNOWLEDGMENTS

We thank Dr. Charis Drummer and Dr. Eva Wolf for providing the marmoset fetal material and Nicole Umland for the histological analysis of teratomas. No specific funding has been received for the work on this manuscript.

## CONFLICT OF INTEREST

The authors declare no conflict of interest.

## Notes

### Competing Interest Statement

The authors have declared no competing interest.

## REFERENCES

Acampora, D., Giovannantonio, L. G. D., Simeone, A. (2013). Otx2 is an intrinsic determinant of the embryonic stem cell state and is required for transition to a stable epiblast stem cell condition. Development 140, 43–55.

Bauer, G., Elsallab, M., Abou-El-Enein, M. (2018). Concise Review: A Comprehensive Analysis of Reported Adverse Events in Patients Receiving Unproven Stem Cell-Based Interventions. Stem Cells Transl. Med. 7, 676–685.

Choi, J., Huebner, A.J., Clement, K., Walsh, R.M., Savol, A., Lin, K., Gu, H., Di Stefano, B., Brumbaugh, J., Kim, S.Y., et al. (2017). Prolonged Mek1/2 suppression impairs the developmental potential of embryonic stem cells. Nature 548, 219–223.

Debowski, K., Warthemann, R., Lentes, J., Salinas-Riester, G., Dressel, R., Langenstroth, D., Gromoll, J., Sasaki, E., and Behr, R. (2015). Non-viral generation of marmoset monkey iPS cells by a six-factor-in-one-vector approach. PLoS One 10: e0118424.

Fleifel, D., Rahmoon, M.A., AlOkda, A., Nasr, M., Elserafy, M., and El-Khamisy, S.F. (2018). Recent advances in stem cells therapy: A focus on cancer, Parkinson’s and Alzheimer’s. J. Genet. Eng. Biotechnol. 16, 427–432.

Itakura, G., Kawabata, S., Ando, M., Nishiyama, Y., Sugai, K., Ozaki, M., Iida, T., Ookubo, T., Kojima, K., Kashiwagi, R., et al. (2017). Fail-safe system against potential tumorigenicity after transplantation of iPSC derivatives. Stem Cell Reports 8, 673–684.

Jho, E.H., Zhang, T., Domon, C., Joo, C.K., Freund, J.N., and Cos-tantini, F. (2002). Wnt/beta-catenin/Tcf signaling induces the tran-scription of Axin2, a negative regulator of the signaling pathway.Mol. Cell. Biol. 22, 1172–1183

Kinney, R.M., Johnson, B.J., Welch, J.B., Tsuchiya, K.R., and Trent, D.W.(1989). The full-length nucleotide sequences of the virulent Trinidad donkey strain of Venezuelan equine encephalitis virus and its attenuated vaccine derivative, strain TC-83. Virology 170, 19–30.

Ma, H., Zhai, J., Wan, H., Jiang, X., Wang, X., Wang, L., Xiang, Y., He, X., Zhao, Z.A., Zhao, B., et at. (2019). In vitro culture of cynomolgus monkey embryos beyond early gastrulation. Science 366, eaax7890.

Martin, U. (2017). Therapeutic Application of Pluripotent Stem Cells: Challenges and Risks. Front. Med. 4:229. eCollection 2017.

Nakajima, M., Yoshimatsu, S., Sato, T., Nakamura, M., Okahara, J., Sasaki, E., Shiozawa, S., and Okano, H. (2019). Establishment of induced pluripotent stem cells from common marmoset fibroblasts by RNA-based reprogramming. Biochem. And Biophys. Res. Commun. 515, 593–599.

Nakamura, M., and Okano, H. (2013). Cell transplantation therapies for spinal cord injury focusing on induced pluripotent stem cells. Cell Res. 23, 70–80.

Niu, Y., Sun, N., Li, C., Lei, Y., Huang, Z., Wu, J., Si, C., Dai, X., Liu, C., Wei, J., et al. (2019). Dissecting primate early post-implantation development using long-term in vitro embryo culture. Science 366, eaaw5754.

Okita, K., Ichisaka, T, and Yamanaka, S. (2007). Generation of germline-competent induced pluripotent stem cells. Nature 448, 313–317.

Okita, K., Matsumura, Y., Sato, Y., Okada, A., Morizane, A., Okamoto, S., Hong, H., Nakagawa, M., Tanabe, K., Tezuka, K.I., et al. (2011). A more efficient method to generate integration-free human iPS cells. Nat. Methods 8, 409–412.

Petrakova, O., Volkova, E., Gorchakov, R., Paessler, S., Kinney, R.M., and Frolov, I. (2005). Noncytopathic replication of Venezuelan equine encephalitis virus and eastern equine encephalitis virus replicons in Mammalian cells. J. Virol. 79, 7597–7608.

Ross, C. N., Davis, K, Dobek, G., and Tardif, S. D. (2012). Aging Phenotypes of Common Marmosets (Callithrix jacchus) Journal of Aging Research Volume 2012, Article ID 567143.

Sasaki, E., Suemizu, H., Shimada, A., Hanazawa, K., Oiwa, R., Kamioka, M., Tomioka, I., Sotomaru, Y., Hirakawa, R., Eto, T., et. al. (2009). Generation of transgenic non-human primates with germline transmission. Nature 459, 523–727.

Sato, K., Oiwa, R., Kumita, W., Henry, R., Sakuma, T., Ito, R., Nozu, R., Inoue, T., Katano, I., Sato, K., et al.. (2016). Cell Stem Cell 19, 127–138.

Schlaeger, T.M., Daheron, L., Brickler, T.R., Entwisle, S., Chan, K., Cianci, A., DeVine, A., Ettenger, A., Fitzgerald, K., and Godfrey, M. (2015). A comparison of non-integrating reprogramming methods. Nat. Biotechnol. 33, 58–63.

Senos, R., Benedicto, H.G., del Rio do Valle, C.M. , del Rio do Valle, R., Nayudu, P.L., Kfoury Junior, J.R., Bombonato, P.P. (2014). Gross morphometry of the heart of the common marmoset. Folia Morphol. 73, 37–41.

Shimozawa, A., Ono, M., Takahara . D., Tarutani, A., Imura, S., Masuda-Suzukake, M., Higuchi, M., Yanai, K., Hisanaga, S., and Hasegawa, M. (2017). Propagation of pathological α-synuclein in marmoset brain. Acta Neuropathol. Commun. 5: 12.

Shroff, G. (2018). A review on stem cell therapy for multiple sclerosis: special focus on human embryonic stem cells. Stem Cells Cloning 11, 1–11.

Solis, M.A., Moreno Velásquez, I., Correa, R., and Huang, L.L.H. (2019). Stem cells as a potential therapy for diabetes mellitus: a call-to-action in Latin America. Diabetol. Metab. Syndr. 11: 20

Song, C. G., Zhang, Y. Z., Wu, H. N., Cao, X. L., Guo, C. J., Li, Y. Q., Zheng, M. H., and Han, H. (2018). Stem cells: a promising candidate to treat neurological disorders Neural Regen. Res. 13, 1294–1304.

Takahashi, K., Tanabe, K., Ohnuki, M., Narita, M., Ichisaka, T., Tomoda, K., and Yamanaka, S. (2007). Induction of pluripotent stem cells from adult human fibroblasts by defined factors. Cell 131, 861–872.

Tardif, S.D., and Ziegler, T.E. (2011). The marmoset as a model of aging and age-related diseases. ILAR J. 52, 54–65.

Tiburcy, M., Hudson, J. E., Balfanz, P., Schlick, S., Meyer, T., Chang Liao, M. L., Levent, E., Raad, F., Zeidler, S., Wingender, E., et al. (2017). Defined engineered human myocardium with advanced maturation for applications in heart failure modeling and repair. Circulation 135, 1832–1847.

Tidball, A.M., Neely, M.D., Chamberlin, R., Aboud, A.A., Kumar, K.K., Han, B., Bryan, M.R., Aschner, M., Ess, K.C., and Bowman, A.B. (2016). Genomic Instability Associated with p53 Knockdown in the Generation of Huntington’s Disease Human Induced Pluripotent Stem Cells. PLoS One 11:e0150372.

Tomioka, I., Maeda, T., Shimada, H., Kawai, K., Okada, Y., Igarashi, H., Oiwa, R., Iwasaki, T., Aoki, M., Kimura, T., et al.. (2010). Generating induced pluripotent stem cells from common marmoset (Callithrix jacchus) fetal liver cells using defined factors, including Lin28 Genes Cells 15, 959–969.

Tomioka, I., Ishibashi, H., Minakawa, E.N., Motohashi, H.H., Takayama, O., Saito, Y., Popiel, H.A., Puentes, S., Owari, K., Nakatani, T., et al. (2017). Transgenic monkey model of the polyglutamine diseases recapitulating progressive neurological symptoms. eNeuro ENEURO 4, 0250–16.

Vermilyea, S.C., Guthrie, S., Meyer, M., Smuga-Otto, K., Braun, K., Howden, S., Thomson, J.A., Zhang, S.C., Emborg, M.E., and Golos, T.G. (2017). Induced pluripotent stem cell-derived dopaminergic neurons from adult common marmoset fibroblasts. Stem Cells Dev. 26, 1225–1235.

Watanabe, T., Yamazaki, S., Yoneda, N., Shinohara, H., Tomioka, I., Higuchi, Y., Yagoto, M., Ema, M., Suemizu, H., Kawai, K., et al. (2019). Highly efficient induction of primate iPS cells by combining RNA transfection and chemical compounds. Genes Cells 24, 473–484.

Wu, Y., Zhang, Y., Mishra, A., Tardif, S.D., and Hornsby, P.J. (2010). Generation of induced pluripotent stem cells from newborn marmoset skin fibroblasts. Stem Cell Res. 4, 180–188.

Yoshioka, N., Gros, E., Li, H.R., Kumar, S., Deacon, D.C., Maron, C., Muotri, A.R., Chi, N.C., Fu, X.D., Yu, B.D., et al. (2013). Efficient generation of human iPSCs by a synthetic self-replicative RNA. Cell Stem Cell 13, 246–254.

Yu, J., Hu, K., Smuga-Otto, K., Tian, S., Stewart, R., Slukvin, I.I., and Thomson, J.A. (2009). Human induced pluripotent stem cells free of vector and transgene sequences. Science 324, 797–801.

Yun, J.W., Ahn, J. B., and Kang, B. C. (2015). Modeling Parkinson’s disease in the common marmoset (Callithrix jacchus): overview of models, methods, and animal care. Lab Anim. Res. 31, 155–165.

Zhou, X., Chadarevian, J.P., Ruiz, B., Ying, Q.L. (2017). Cytoplasmic and Nuclear TAZ Exert Distinct Functions in Regulating Primed Pluripotency. Stem Cell Reports 9,732–741.

