## Supplementary Figure S1 for "Controlling the switch from neurogenesis to pluripotency during marmoset monkey somatic cell reprogramming with self-replicating mRNAs and small molecules"

### Supplemental Information

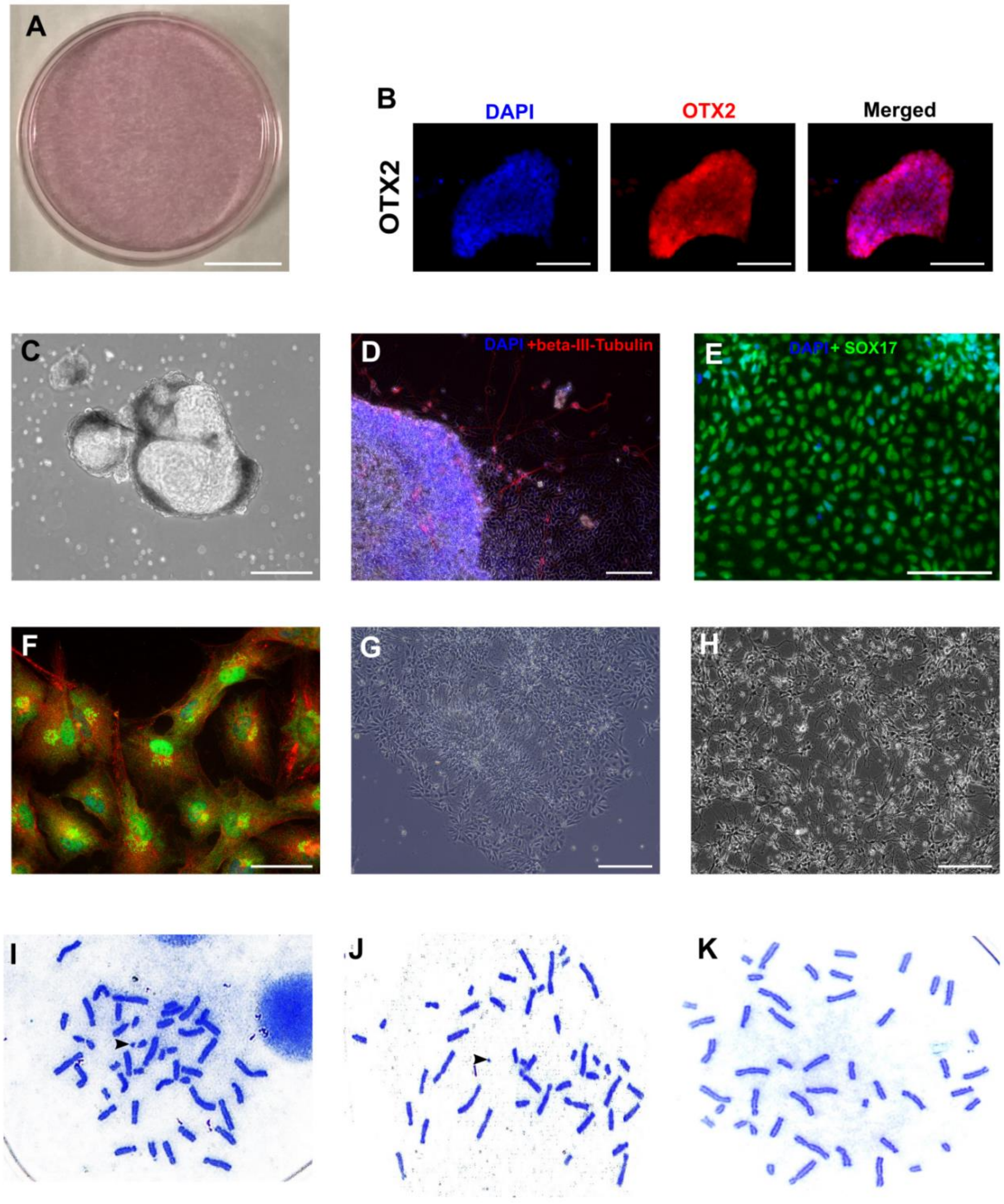


**Figure S1. Characterization of marmoset iPSCs: marker expression, *in vitro* differentiation, and karyotyping. A)** Alkaline phosphatase activity in iPSC culture. **B)** OTX2 immunofluorescence in marmoset iPSC colony. **C)** Embryoid body in suspension. **D)** Embryoid body outgrowth on glass coverslip immunostained with DAPI (blue) and anti-beta-III-Tubulin antibody (red fluorescence). **E)** Embryoid body outgrowth on glass coverslip immunostained with DAPI (blue) and anti-SOX17 (green fluorescence). **F-H)** Directed neural differentiation of marmoset iPSCs: **F**) Marmoset iPSC-derived NSC-like cells expressing PAX6 (green fluorescence) and NESTIN (red fluorescence). **G)** Neural rosette formed after culture on poly-L-ornithine and laminin-coated surface. **H)** Neurons at the end of differentiation. **I-K)** Representative karyotypes of two male and one female iPSC lines with normal chromosome numbers (46 XY/XX). The Y chromosomes are indicated with arrowheads. (Scale bars: A = 2 cm; B, D, E = 100 µm; C = 200 µm; F = 50 µm; G and H = 200 µm).
